# Natural interploidy hybridization among the key taxa involved in the origin of horticultural chrysanthemums

**DOI:** 10.1101/2021.07.29.454281

**Authors:** Shuai Qi, Alex D. Twyford, Junyi Ding, James S. Borrell, Yueping Ma, Nian Wang

**Affiliations:** State Forestry and Grassland Administration Key Laboratory of Silviculture in Downstream Areas of the Yellow River, College of Forestry, Shandong Agricultural University, Tai’an 271018, China; Institute of Evolutionary Biology, Ashworth Laboratories, the University of Edinburgh, Edinburgh EH9 3FL, UK; Royal Botanic Garden Edinburgh, 20A Inverleith Row, Edinburgh EH3 5LR, UK; Royal Botanic Gardens Kew, Richmond, Surrey TW9 3AB, UK; College of Life and Health Sciences, Northeastern University, Shenyang 110004, China; Mountain Tai Forest Ecosystem Research Station of State Forestry and Grassland Administration, College of Forestry, Shandong Agricultural University, Tai’an 271018, China; State Key Laboratory of Crop Biology, Shandong Agricultural University, Tai’an, China

**Author notes:** Author for correspondence: Nian Wang.

**Keywords:** *Chrysanthemum*, hybrid, microsatellite marker, symmetrical introgression, *trn*L-*trn*F

## Abstract

Understanding hybridization and introgression between natural plant populations can give important insights into the origins of cultivated species. Recent studies suggest differences in ploidy may not create such strong reproductive barriers as once thought, and thus studies into cultivated origins should examine all co-occurring taxa, including those with contrasting ploidy levels. Here, we characterized hybridization between *Chrysanthemum indicum, Chrysanthemum vestitum* and *Chrysanthemum vestitum* var. *latifolium*, the most important wild species involved in the origins of cultivated chrysanthemums. We analysed population structure of 317 *Chrysanthemum* accessions based on 13 microsatellite markers and sequenced chloroplast *trn*L-*trn*F for a subset of 103 Chrysanthemum accessions. We identified three distinct genetic clusters, corresponding to the three taxa. We detected 20 hybrids between species of different ploidy levels, of which 19 were between *C. indicum* (4x) and *C. vestitum* (6x) and one was between *C. indicum* and *C. vestitum* var. *latifolium* (6x). Fourteen hybrids between *C. indicum* and *C. vestitum* were from one of the five study sites. *Chrysanthemum vestitum* and *C. vestitum* var. *latifolium* share only one chloroplast haplotype. The substantially different number of hybrids between hybridizing species was likely due to different levels of reproductive isolation coupled with environmental selection against hybrids. In addition, human activities may play a role in the different patterns of hybridization among populations.

## 1. Introduction

Hybridization has played an important role in plant domestication and diversification through human history (Arnold, 2014; Heslop-Harrison and Schwarzacher, 2007; Cornille et al., 2014). Multiple important crops have been generated through hybridization either between wild species or through introgression from crop wild relatives into cultivated lineages. Major examples include modern strawberries (*Fragaria ananassa*) (Bringhurst and Voth, 1984) and triploid bananas (Simmonds and Shepherd, 1955; Heslop-Harrison and Schwarzacher, 2007), but also ornamental species such as tree peonies (*Paeonia suffruticosa*) (Zhou et al., 2014), cherry (*Prunus yedoensis*) (Baek et al., 2018) and dahlia (*Dahlia variabilis*) (Saar et al., 2003). Understanding the frequency, phylogenetic distribution and propensity for hybridization in wild populations could not only inform breeders as to the possible range of interspecific hybrids, but could also reduce the laborious, time consuming and frequently unsuccessful process of artificial crossing and *de novo* hybrid generation (Lim et al., 2008; Kuligowska et al., 2016).

Hybridization occurs more easily between species of the same ploidy level than differing ploidy levels. For example, hybridization between diploid and tetraploid species is often limited as triploid hybrids are usually inviable and less fit, preventing backcross formation (Wang et al., 2014; Zohren et al., 2016; Husband and Sabara, 2003). However, species of contrasting higher ploidy levels appear to have weaker reproductive barriers and hybridize more easily than diploids and tetraploids (Sonnleitner et al., 2013; Sutherland and Galloway, 2017). For example, within the *Campanula rotundifolia* polyploid complex, postzygotic isolation was lower in tetraploid–hexaploid species than in diploid–tetraploid crosses (Sutherland and Galloway, 2017). To date only a small number of studies have investigated hybridization between higher ploidy levels, and the prevalence of higher level cross-ploidy hybridization across plant families remains unclear.

*Chrysanthemum* L. (Asteraceae) provides an excellent model for studying hybridization between high ploidy levels, with ploidy ranging from diploid (2n = 2x = 18) to decaploid (2n = 10x = 90) (Ma et al., 2015; Zhou and Wang, 1997; Tahara, 1915; Li et al., 2013; Luo et al., 2017) and with different cytotypes within species (Chen, 2012; Yan et al., 2019; Dowrick, 1952; Dowrick, 1953; Liu et al., 2012). *Chrysanthemum* includes approximately 37 wild species, of which 17 occur in China (Shih and Fu, 1983) where they have captured great public interest. Multiple wild species have been crossed by humans to generate numerous cultivars, and they are among the most famous Chinese flowers, with significant commercial and medicinal value (Kim and Lee, 2005; Shahrajabian et al., 2019). The evolutionary history of polyploidy within *Chrysanthemum* is currently unknown, though some polyploids are thought to be allopolyploid in origin (Liu et al., 2012; Chen et al., 1996) and subject to multiple historical polyploidization events (Yang et al., 2006).

Chrysanthemums were first cultivated in China ∼1600 years ago, and were later introduced to Japan and Europe (Chen, 2012; Shih et al., 2011; Chen, 1985). Modern cultivated chrysanthemums are mainly hexaploids and hybridization and subsequent artificial selection are thought to give rise to numerous cultivars (Chen, 1985; Dai et al., 2002). The ancestry of modern chrysanthemums remains elusive, but several wild species are thought to be involved, including *C. indicum* (4x), *C. vestitum* (6x), *C. lavandulifolium* (2x), *C. nankingense* (2x) and *C. zawadskii* (2x) (Chen, 1985; Dai and Chen, 1997; Fukai, 2003; Ma et al., 2016; Ma et al., 2020). *Chrysanthemum indicum* and *C. vestitum* are key species in the origin and evolution of cultivated chrysanthemums (Dai et al., 2002; Dai et al., 1998) based on two lines of evidence. First, ancient literature documents multiple uses for *C. indicum* in central China, which is consistent with the geographic origin of modern cultivars (see Chen, 2012). Second, artificial hybridization between *C. indicum* and *C. vestitum* can generate hybrids resembling the prototype of modern chrysanthemums (Chen, 2012).

In this study, we investigate whether hybridization occurs between *Chrysanthemum* species with different ploidy levels, focusing on *C. indicum* and *C. vestitum* as well as varieties of those species. We ask: (1) Does interploidy hybridization naturally occur between tetraploid and hexaploid *Chrysanthemum* species, which are likely involved in origin of modern cultivated chrysanthemums? (2) If hybridization occurs, is there evidence for a greater propensity at high ploidy levels? (3) Do some modern chrysanthemum cultivars share chloroplast haplotypes with the three wild *Chrysanthemum* taxa? To this end, we genotyped 317 samples at 13 microsatellite markers and sequenced chloroplast *trn*L-*trn*F for a subset of 103 samples. In addition, we extracted *trn*L-*trn*F from the chloroplast genomes of 28 taxa, representing 15 wild species of *Chrysanthemum*, 12 cultivars and one sample of *Ajania varifolium*. We analyze these data in a population genetic and phylogeography context, and use them to investigate a case where traditionally reproductive isolation caused by differences in ploidy levels would be expected to be strong.

## 2. Material and Methods

### 2.1 Study species

*Chrysanthemum indicum* is tetraploid with a wide distribution across China, though narrowly distributed diploid and hexaploid cytotypes have also been reported (Liu et al., 2012). *Chrysanthemum vestitum* is a hexaploid distributed across Hubei and Henan provinces in central China (Zhao and Chen, 1999) though the variety *C. vestitum* var. *latifolium* is hexaploid with a restricted distribution in the Dabie Mountains in Anhui province.

The three taxa are outcrossing perennials (Chen, 2012) and are likely to be pollinated by bees (personal observations, N. Wang). Morphologically, *C. indicum* has yellow florets and smooth leaves with deep serrations. Both *C. vestitum* and *C. vestitum* var. *latifolium* have white floret and pubescent leaves and stems (Fig. 1). Putative hybrids either with an intermediate morphology or ploidy level between *C. indicum* and *C. vestitum* have been found in localities where the species co-occur (Nakata et al., 1992; Zhao and Chen, 1999). These hybrids exhibit continuous phenotypes between *C. indicum* and *C. vestitum* (Zhao and Chen, 1999), indicating the existence of putative hybrid swarms.

**Fig. 1.**
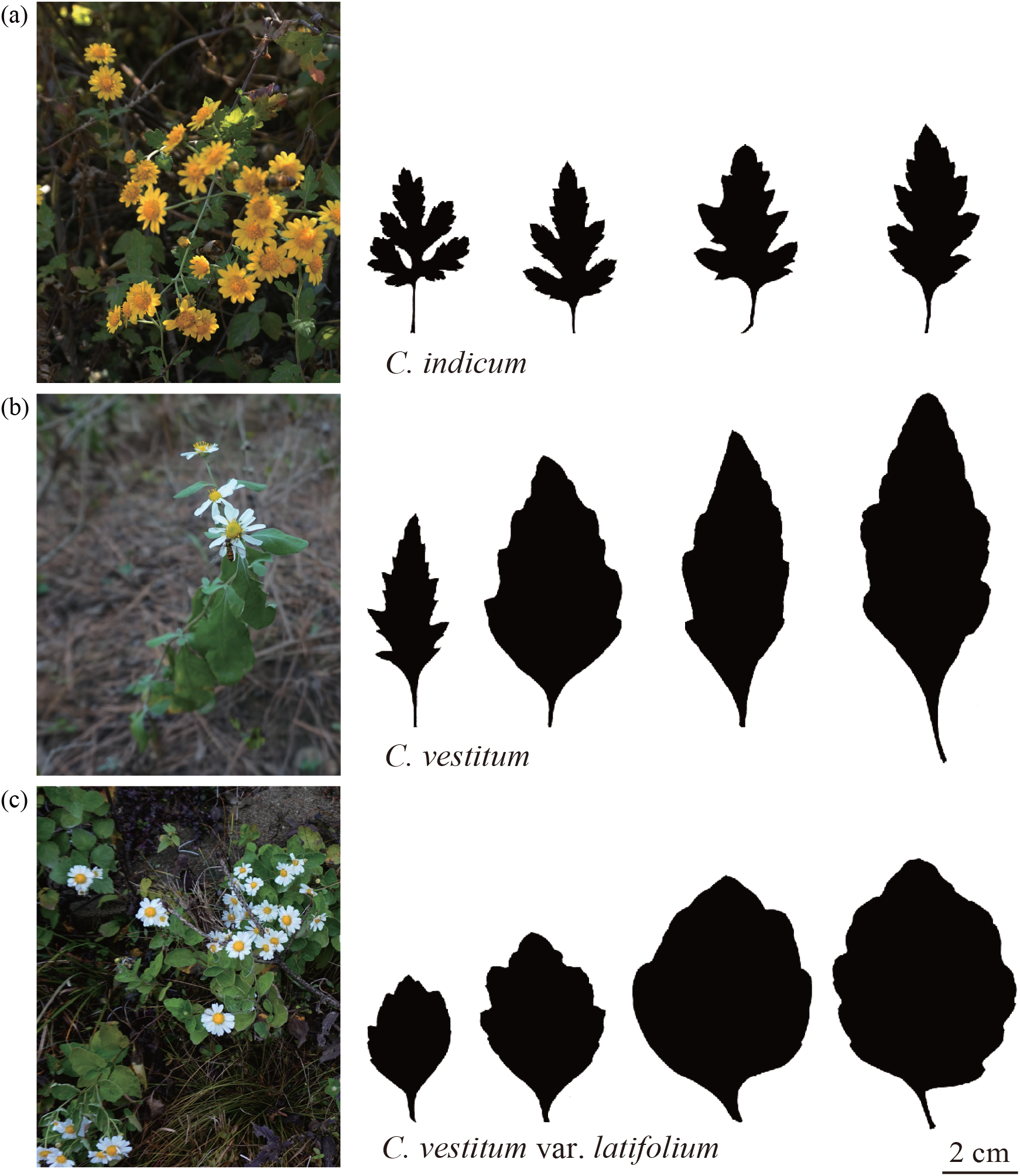
Diversity present in wild *Chrysanthemum* species included in this study. Individual photographs and leaf morphology for *C. indicum* (a), *C. vestitum* (b) and *C. vestitum* var. *latifolium* (c).

### 2.2 Sampling across hybridizing populations

To understand the extent and distribution of hybridization, we surveyed sympatric populations of the three focal taxa: *C. indicum, C. vestitum*, and *C. vestitum* var. *latifolium*. We identified and sampled five populations where *C. indicum* and *C. vestitum* co-occur and two where *C. indicum* and *C. vestitum* var. *latifolium* co-occur (Table S1). In addition, we collected *C. indicum* from two allopatric populations, TA and ZP (Table S1). Samples were collected at random within populations, ensuring at least ten meter spacing between individuals. Healthy and pest free leaf tissue was collected and stored in silica gel. A total of 317 samples were collected including between 14 and 74 from each of the seven hybridizing populations, five from TA and three from ZP (Table S1). A Global Position System (GPS, Unistrong) was used to record the coordinates of each population. Sampling locations are illustrated in Fig. 2.

**Fig. 2.**
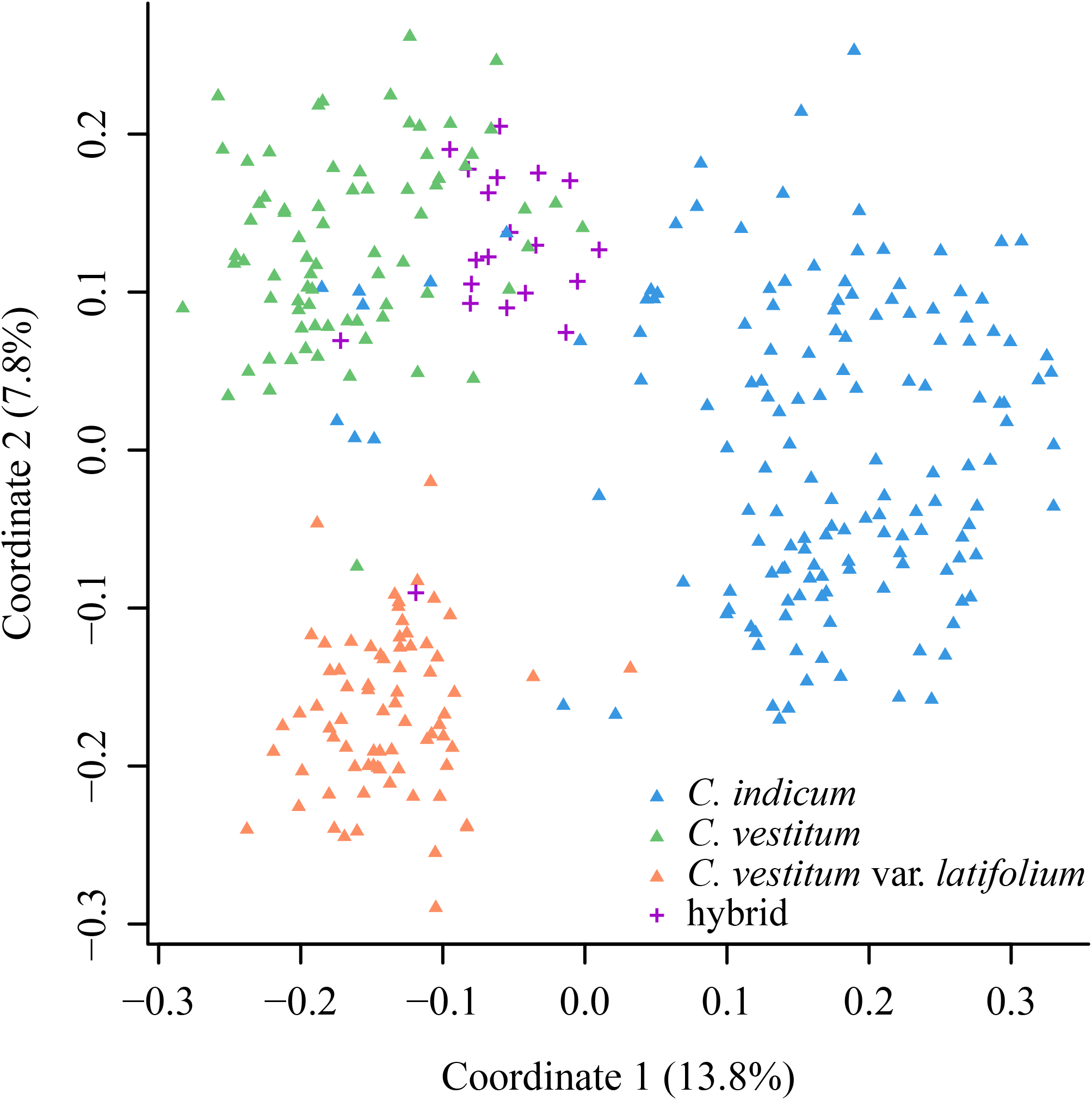
Principal coordinate analysis of *Chrysanthemum* samples based on 13 microsatellite markers. Hybrids were defined based on Q scores in a STRUCTURE analysis (see text).

### 2.3 Microsatellite genotyping

Genomic DNA was isolated from dried leaves of all individuals, following a modified 2x CTAB (cetyltrimethylammoniumbromide) protocol (Wang et al., 2013). The quality of isolated genomic DNA was assessed on 1.0 % agarose gels, and then diluted to a concentration of 10-20 ng/ul for genotyping and sequencing. Thirteen microsatellite loci were used for genotyping (Zhang et al., 2014; Jo et al., 2015). The 5’ terminus of the forward primer was labeled with FAM, HEX or TAM fluorescent probes. Each microsatellite locus was amplified individually prior to being combined into four multiplexes (Table S2). The PCR protocol follows Hu et al. (2019).

### 2.4 Population genetic analysis

It is often difficult to assign microsatellite genotypes for mixed ploidy species, as the frequency of different alleles can be difficult to quantify. In hexaploids, each microsatellite locus would be expected to have up to six alleles per individual. We chose to score each allele separately using the software GENEMARKER 2.4.0 (Softgenetics), and checked each genotype manually. We then calculated allele richness for each population using FSTAT 2.9.4 (Goudet, 1995) and performed principal coordinate (PCO) analysis in POLYSAT 1.7-4 (Clark and Jasieniuk, 2011), based on Bruvo’s pairwise genetic distances (Bruvo et al., 2004).

We performed STRUCTURE analysis for each hybridizing population separately using STRUCTURE 2.3.4 (Pritchard et al., 2000) with ploidy specified as 6n. We combined XG, NX and PH into one hybridizing population as the three localities are separated by only a few kilometers. We set the number of genetic clusters (K) to 2 when analyzing the genetic structure of each hybridizing population as only two parental species are present. The allopatric *C. indicum* populations (TA and ZP) were used as a reference population.

In addition, to identify the most likely K value across populations we included all populations in a combined STRUCTURE analysis, testing K values from 1 to 10. The number of genetic clusters was estimated using the Evanno test (Evanno et al., 2005) in the program Structure Harvester 0.6.94 (Earl and vonHoldt, 2012). Ten replicates of the STRUCTURE analysis were performed with 1,000,000 iterations and a burn-in of 100,000 for each run. The admixture model, with an assumption of correlated allele frequencies, was used. Individuals were assigned to clusters based on the highest membership coefficient averaged over the ten independent runs. Replicate runs were grouped based on a symmetrical similarity coefficient of >0.9 using the Greedy algorithm in CLUMPP 1.1.2 (Jakobsson and Rosenberg, 2007) and visualized in DISTRUCT 1.1 (Rosenberg, 2004). In populations where *C. indicum* and *C. vestitum* or *C. indicum* and *C. vestitum* var. *latifolium* co-occur, we estimated Q scores in STRUCTURE with 95% confidence intervals to define pure *C. indicum*, pure *C. vestitum*, pure *C. vestitum* var. *latifolium* or putative hybrids. We distinguish individuals with confidence intervals overlapping 1 as pure *C. indicum*, with 0 as pure *C. vestitum* and those remaining as putative hybrids, in populations where the two species co-occur. Similarly, in populations where *C. indicum* and *C. vestitum* var. *latifolium* co-occur, we distinguish individuals with confidence intervals overlapping with 1 as pure *C. indicum*, with 0 as pure *C. vestitum* var. *latifolium* and those remaining as hybrids.

We compared the average allele number per individual between *C. indicum, C. vestitum, C. vestitum* var. *latifolium* and their hybrids at each microsatellite locus, using the Kruskal–Wallis test in the R package agricolae v1.3-3 (de Mendiburu, 2020). We would expect that the average allele number is higher for *C. vestitum* and *C. vestitum* var. *latifolium* than *C. indicum*. In addition, we tested if introgression was symmetric between *C. indicum* and *C. vestitum* or between *C. indicum* and *C*. *vestitum* var. *latifolium* in each hybridizing population, using the function =wilcox.test’ in R v4.0.3 (R Core Team, 2020). The putative hybrids identified as above excluded from such comparisons.

### 2.6 *trn*L-*trn*F sequencing and analysis

In order to detect the potential maternal parents of the hybrids, and to estimate the haplotype diversity of these taxa, we amplified *trn*L-*trn*F for a subset of 103 samples, including between five and 34 individuals from each population. Reactions were performed in 20 ul volumes containing 13 ul ddH_2_O, 5 ul 2×Taq PCR mix (TIANGEN, China), 0.5 ul each primer (*trn*L and *trn*F (Taberlet et al., 1991)) and 1 ul DNA template. PCR products were outsourced for purification and sequencing, to Qingdao, China. Sequences were manually edited and aligned using BioEdit v7.2.5 (Hall, 1999). The R package pegas v 0.14 (Paradis, 2010) was used to construct haplotype networks, using default settings, with gaps treated as a fifth state. The total number of sites, polymorphic sites, parsimony informative sites, and nucleotide diversity were computed using DnaSPv6.12.03 (Rozas et al., 2017). All sequences obtained in this study were submitted to GenBank with accession number MZ032043 -MZ032145.

To aid in a broader phylogenetic analysis, we extracted the *trn*L-*trn*F region from the available whole chloroplast genomes for 15 of the 17 wild species of *Chrysanthemum* occurring in China, 12 cultivars and *Ajania varifolium*. A phylogenetic tree was estimated using the maximum-likelihood method (ML) in RAxML v. 8.1.16 (Stamatakis, 2006). *Ajania varifolium* was selected as the outgroup. A rapid bootstrap analysis with 100 bootstrap replicates and 10 tree searches was performed under the GTR + GAMMA nucleotide substitution model. The consensus tree generated from the bootstrap replicates was visualized in FigTree v.1.3.1 (Rambaut and Drummond, 2009).

## 3 Results

### 3.1 Hybridization across ploidy levels inferred from microsatellites

Genetic diversity estimates were similar among the three taxa and hybrids, with allelic richness ranging from 3.74 to 4.26 and gene diversity from 0.75 to 0.84 (Table S3). On average, the number of alleles scored in *C. vestitum* and *C. vestitum* var. *latifolium* was significantly higher than *C. indicum* at eight and seven loci, respectively (P < 0.05); this was expected for a hexaploid possessing more chromosome copies than a tetraploid species (Fig. S1).

Principal Coordinate (PCO) analysis based on Bruvo’s genetic distances among all samples revealed three clusters, with coordinates 1 and 2 explaining 13.8% and 7.8% of the total variation, respectively (Fig. 2). Coordinate 1 separated *C. indicum* from *C. vestitum* and *C. vestitum* var. *latifolium* and coordinate 2 separated *C. vestitum* from *C. vestitum* var. *latifolium* (Fig. 2). Most hybrids identified by the STRUCTURE analysis fell between *C. indicum* and *C. vestitum* in the PCO plot (Fig. 2).

The combined STRUCTURE analysis across all samples identified K = 3 as the optimal K value (Fig. S2a), with the three clusters corresponding to *C. indicum, C. vestitum* and *C. vestitum* var. *latifolium* (Fig. 3a). At K = 2, *C. indicum* formed one cluster and *C. vestitum* and *C. vestitum* var. *latifolium* formed another cluster (Fig. S2b), supporting the close relationship between these two taxa and the inference that *C. vestitum* var. *latifolium* is a subspecies of *C. vestitum*. Interestingly, there are three *C. vestitum* individuals possessing a considerable level of introgression from *C. vestitum* var. *latifolium* and one *C. indicum* individual in population TZ with substantial introgression from *C. vestitum* (Fig. 3a).

**Fig. 3.**
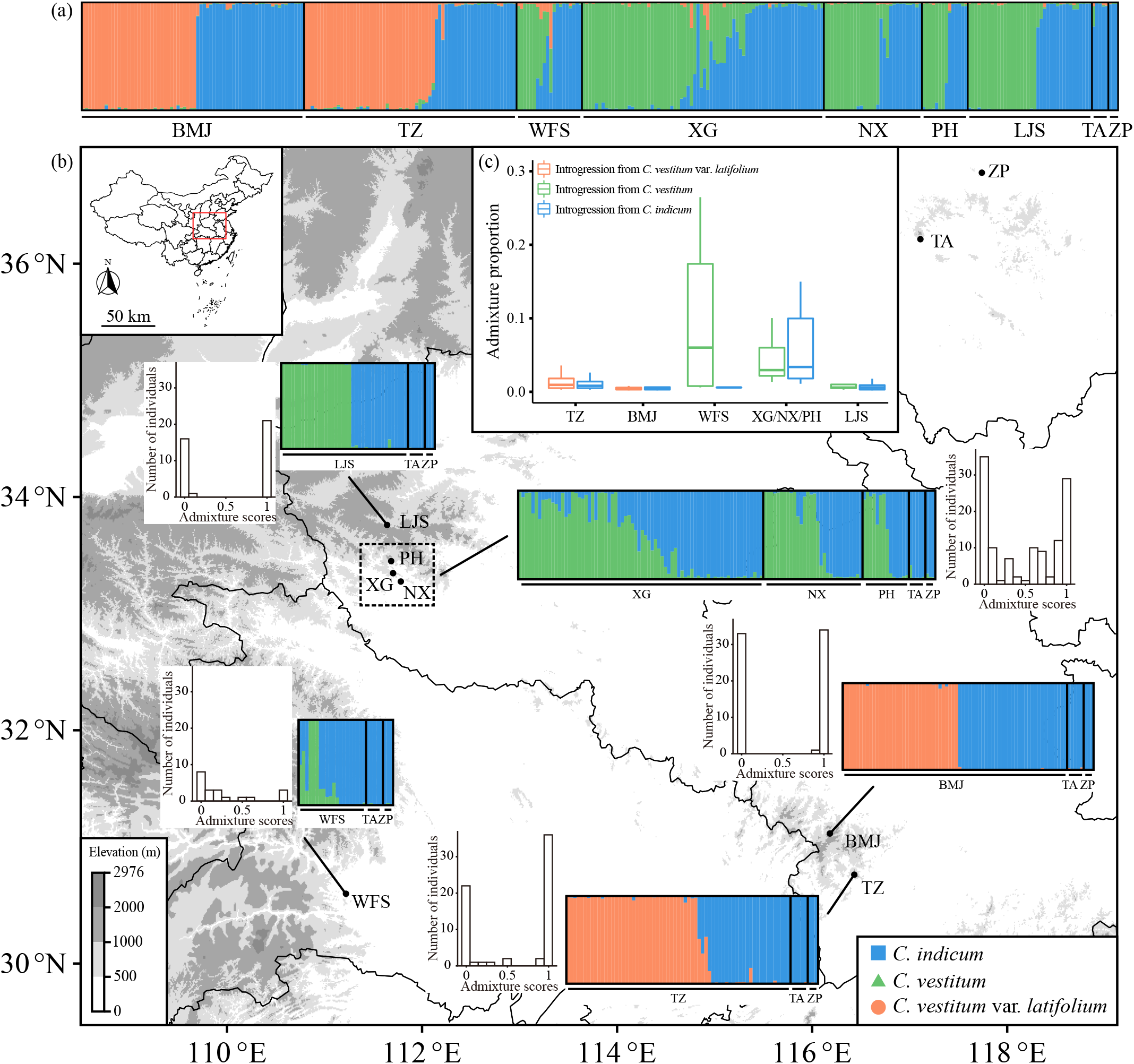
Hybridization among *C. indicum, C. vestitum* and *C. vestitum* var. *latifolium*. (a) Hybridzation across natural populations in a combined analysis of *C. indicum, C. vestitum* and *C. vestitum* var. *latifolium* with K = 3. (b) Admixture in sympatric population of *C. indicum* and *C. vestitum* and each sympatric population of *C. indicum* and *C. vestitum* var. *latifolium*. Barplots on the left or the right of STRUCTURE plot represent the number of individuals having different levels of genetic admixture. Populations XG, NX and PH were analyzed together due to their short geographic distance. Allopatric populations TA and ZP were served as a control. Blue, green and orange represent *C. indicum, C. vestitum* and *C. vestitum* var. *latifolium*, respectively. (c) Admixture value for each population among *C. indicum, C. vestitum* and *C. vestitum* var. *latifolium*. Hybrids were excluded for such comparisons.

A total of 20 hybrids were detected when STRUCTURE was performed for each hybridizing population separately, of which 19 were hybrids between *C. indicum* and *C. vestitum* and one was a hybrid between *C. indicum* and *C. vestitum* var. *latifolium*. Fourteen hybrids between *C. indicum* and *C. vestitum* were from population XG, two from each of NX and PH and one from WFS. The only hybrid between *C. indicum* and *C. vestitum* var. *latifolium* was from population TZ. No hybrids were detected from population LJS where *C. indicum* and *C. vestitum* co-occur and from BMJ where *C. indicum* and *C. vestitum* var. *latifolium* co-occur (Fig. 3b).

Introgression occurred symmetrically between *C. indicum* and *C. vestitum* and between *C. indicum* and *C. vestitum* var. *latifolium* in all hybridizing populations except WFS where only three *C. indicum* were collected (Fig. 3c). Introgression was limited in populations TZ, BMJ and LJS but was extensive in population XG (Fig. 3c).

### 3.2 Plastid diversity and directionality of hybrid formation

A total of 17 chloroplast haplotypes were detected across samples, with 12, nine, and four in *C. indicum, C. vestitum* and *C. vestitum* var. *latifolium*, respectively (Fig. 4; Table S4). Both *C. indicum* and *C. vestitum* harbored three private haplotypes and *C. vestitum* var. *latifolium* harbored one (haplotype H3). *Chrysanthemum indicum* shared six haplotypes with *C. vestitum* and three haplotypes with *C. vestitum* var. *latifolium*. By contrast, *C. vestitum* only shared the most common haplotype, H6, with *C. vestitum* var. *latifolium*. Frequencies of particular haplotypes varied considerably between species as some haplotypes found in *C. indicum* or *C. vestitum* were absent in *C. vestitum* var. *latifolium* (Fig. 4). Three haplotypes (H6, H16 and H17) were found in hybrids, with haplotype H6 shared among the three taxa, and haplotype H16 found to be private to the hybrid (Fig. 4). One hybrid shared haplotype H17 with *C. indicum*, whereas this haplotype is absent from all *C. vestitum* and *C. vestitum* var. *latifolium* samples (Fig. 4).

**Fig. 4.**
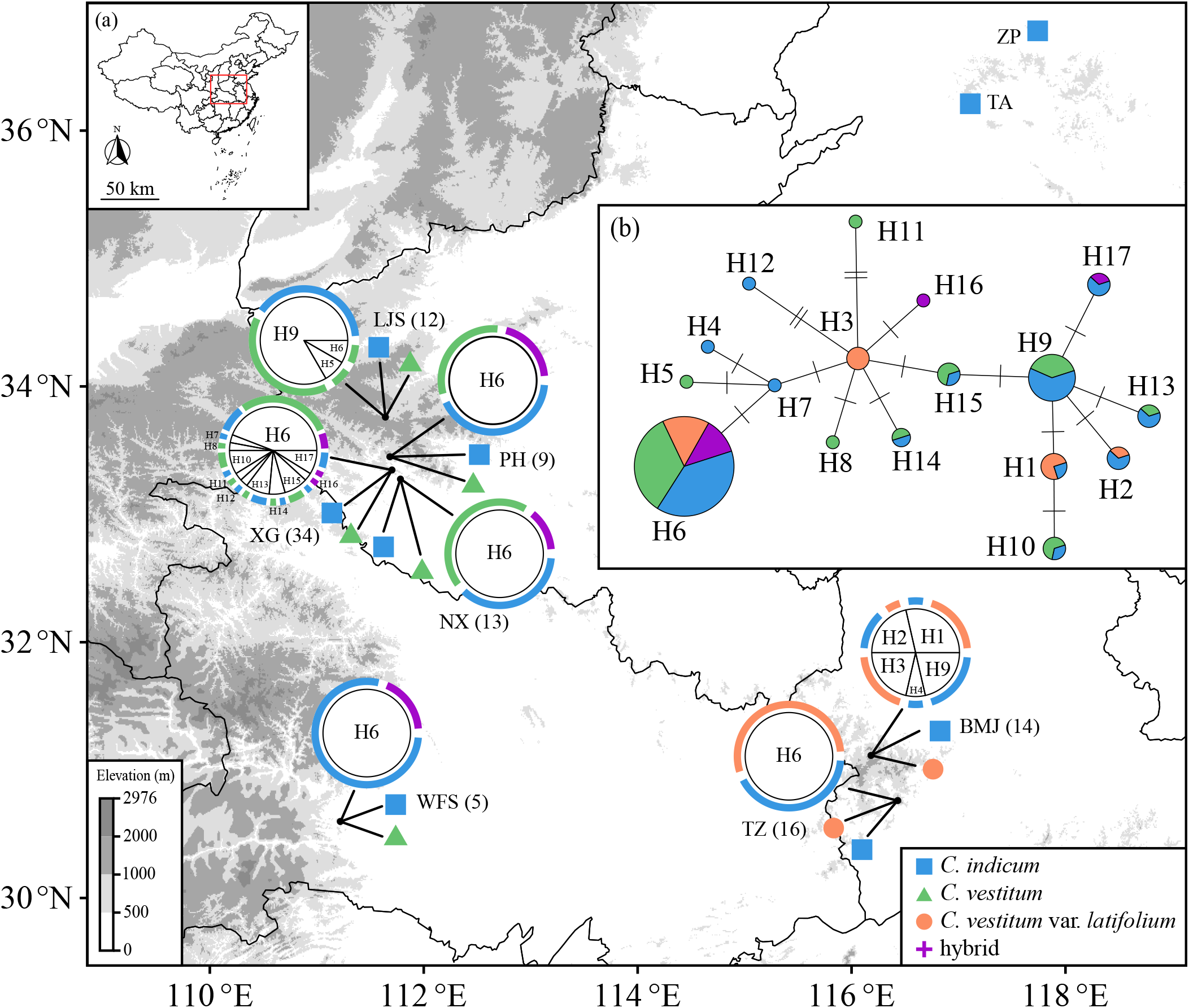
Haplotype network based on plastid *trn*L-*trn*F sequences in natural *Chrysanthemum* populations. (a) The geographic distribution of haplotypes. Number in brackets indicates the number of samples used for sequencing; (b) Haplotype network graph. Each haplotype is represented by a circle with size proportional to the number of individuals. Color within the circle represents species sharing the haplotype.

All modern *C. morifolium* cultivars formed a clade with full support, and this clade was nested in a large monophyletic clade including *C. dichrum, C. zawadskii, C. chanetii* (Fig. 5). Five haplotypes (1, 3, 6, 16 and 17) from the three taxa and hybrid were also nested in this monophyletic clade (Fig. 5).

**Fig. 5.**
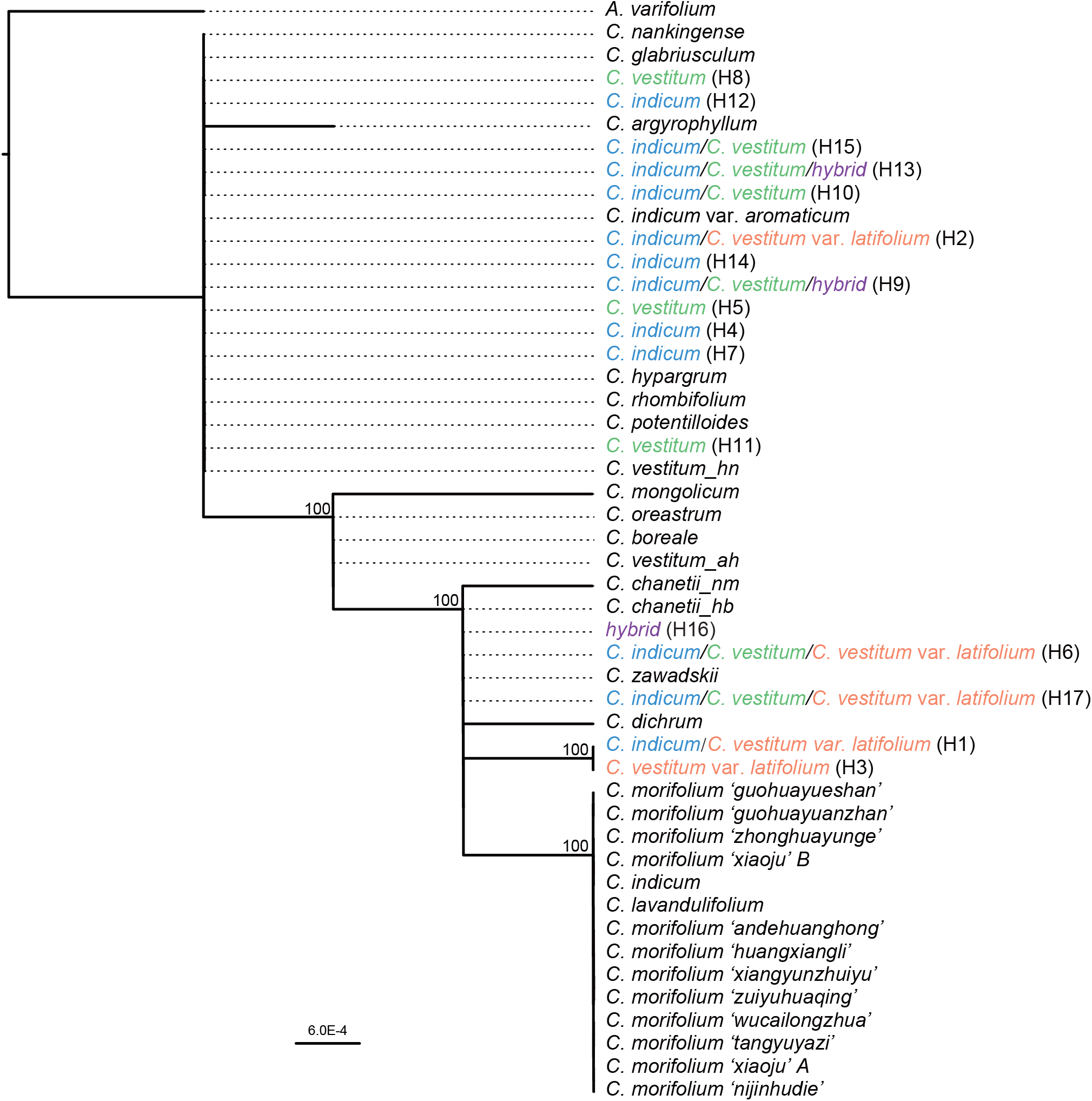
The phylogenetic relationships of *Chrysanthemum* species inferred from the chloroplast *trn*L-*trn*F region. Branch support values above 50% are depicted. Haplotypes identified in the present study are indicated in brackets. Species names before the identified haplotypes represent species sharing the haplotype. Colored in blue, green, orange and purple represent *C. indicum, C. vestitum, C. vestitum* var. *latifolium* and hybrids, respectively.

## 4 Discussion

In this study, we provide genetic evidence of natural interploidy hybridization between two pairs of taxa involved in the formation of modern *Chrysanthemum* horticultural hybrids. We detected more hybrids between *C. indicum* and *C. vestitum* than between *C. indicum* and *C. vestitum* var. *latifolium*, possibly due to different levels of reproductive isolation. In addition, we show that *C. vestitum* var. *latifolium* formed a genetic cluster distinct from *C. vestitum*, and as such deserves its varietal status. Here, we first discuss the importance of hybridization between different ploidy levels in natural *Chrysanthemum* populations, before considering the dynamics of different hybrid swarms. We finish with the wider implications of our findings for understanding the origin of horticultural chrysanthemum hybrids.

### 4.1 Tetraploid-hexaploid hybridization and symmetrical introgression

Interploidy hybridization is more common in genera containing many polyploids, and where species readily co-occur and hybridize, such as *Betula* (Hu et al., 2019; Zohren et al., 2016) and *Spartina* (Ainouche et al., 2003). Diploid-tetraploid hybridization produces mostly sterile triploids, though where hybrids are fertile introgression usually occurs preferentially from diploids to tetraploids (Moraes et al., 2013; Pinheiro et al., 2010). In contrast, hybridization between tetraploids and hexaploids has been proposed to be easier, as pentaploid hybrids are formed frequently (Hülber et al., 2015) and are more fertile than triploids (Sutherland and Galloway, 2017). Consistent with this, we detected 20 hybrids among *C. indicum, C. vestitum* and *C*. *vestitum* var. *latifolium*, indicating incomplete reproductive isolation. The number of hybrids is likely to be underestimated in our study, because of the stringent confidence threshold applied in the STRUCTURE analysis. Three individuals with admixture between 26.2%-31.4 from population XG and two individuals with admixture between 34.5%-48.7% were not supported by Q scores with 95% CIs, but may prove to be hybrids such as later generation backcrosses.

Nineteen out of the 20 hybrids were between *C. indicum* and *C. vestitum* and one was between *C. indicum* and *C. vestitum* var. *latifolium*. This difference may reflect different levels of reproductive isolation. Higher fruit set in artificial *C. indicum*-*C. vestitum* crosses than artificial *C. indicum*-*C. vestitum* var. *latifolium* crosses, partially supports this hypothesis (Zhou, 2009). However, environmental selection against hybrids between *C. indicum* and *C. vestitum* var. *latifolium* may also account for its rarity. *Chrysanthemum indicum* and *C. vestitum* occupy similar habitats and are usually intermixed in sympatric populations (Shuai Qi, personal observations). This may enhance their opportunity for hybridization, while their similar habitat preferences may reduce the chance of ecological selection on the hybrids. In contrast, *C. vestitum* var. *latifolium* and *C. indicum* are adapted to different conditions, and hybrids may fail to survive due to the breakdown of suites of co-adapted genes. Within some diploid-tetraploid systems, introgression is more common from diploids to tetraploids (Zohren et al., 2016; Wang et al., 2020). However, we observe symmetrical introgression between *C. indicum* and *C. vestitum* and between *C. indicum* and *C. vestitum* var. *latifolium*, indicating that hybrids can backcross with both parents. This is in line with recent studies showing that pentaploids can mediate gene flow between species with different ploidy levels (Peskoller et al., 2021). We couldn’t distinguish clearly between male and female parental taxa because *C. indicum* and *C. vestitum* share some plastid haplotypes. However, *C. indicum* can serve as maternal parent as one hybrid shared a haplotype with *C. indicum* whereas this haplotype is absent from all *C. vestitum* and *C. vestitum* var. *latifolium* samples. In addition, one hybrid had a unique haplotype (H16), which is possibly from unsampled *C. indicum* and *C. vestitum* or introgressed from other *Chrysanthemum* species.

### 4.2 Variable hybridizing among populations

The number of hybrids between *C. indicum* and *C. vestitum* varied substantially among the hybridizing populations (Fig. 3b). Fourteen out of 19 hybrids are in population XG, and no hybrids are found in population LJS. These populations are approximately 50 km apart, and differential reproductive isolation seems unlikely to account for such differences. However, we note that population LJS is closer to human habitation and human activities may have an impact on the persistence of hybrids.

Unexpectedly, one hybrid between *C. indicum* and *C. vestitum* var. *latifolium* is from population TZ and none are from BMJ. Moreover, the extent of genetic admixture in TZ seems to be higher than in BMJ (Fig. S1). In TZ, *C. indicum* and *C. vestitum* var. *latifolium* grow closely together, meaning there are enhanced opportunities for hybridization, and likely relaxed selection against hybrids. However, in BMJ, *C. indicum* and *C. vestitum* var. *latifolium* are segregated by altitude; this may limit the survival of hybrids due to environmental selection.

However, under future climate change, *C. indicum* may move to higher altitudes and come into closer contact with *C. vestitum* var. *latifolium*, producing more hybrids, as seen in population TZ. This has implications for conserving *C. vestitum* var. *latifolium. Chrysanthemum indicum* is widespread and abundant whereas *C. vestitum* var. *latifolium* is restricted to the Dabie Mountains. Hybridization between abundant *C. indicum* and rare *C. vestitum* var. *latifolium* may be predicted to drive the rare species to extinction through genetic or demographic swamping (Todesco et al., 2016).

### 4.3 The presence of orphan hybrids

Hybrids usually occur in sympatry with their parental species, but sometimes they occur separately, due to (often human-mediated) long-distance dispersal of hybrids, or natural colonization of sterile hybrid taxa (James and Abbott, 2005). Alternatively parental species may die out due to genetic swamping or competitive exclusion (Huxel, 1999; Levin et al., 1996). Regardless of the mechanism, these result in orphan hybrids (Marques et al., 2010; Groh et al., 2019).

We detected a few individuals showing considerable admixture between *C. vestitum* var. *latifolium* and *C. vestitum* in populations WFS and NX where *C. vestitum* var. *latifolium* is not known to occur. We also detected one *C. indicum* individual showing considerable admixture from *C. vestitum* in population TS in Shandong province. This indicates the presence of hybrids in the absence of one or both parental species, which has been demonstrated in some plant species, such as oaks (Dodd and Afzal-Rafii, 2004) and pines (Lanner and Phillips, 1992). A plausible explanation is the existence of undetected *C. vestitum* var. *latifolium* near populations WFS and NX, or within travelling distance of pollinators or seed dispersal.

### 4.4 Implications for the origins of cultivated chrysanthemums

Our results based on an analysis of plastid *trn*L-*trn*F showed a monophyletic clade composed of *C. lavandulifolium, C. chanetii, C. zawadskii* and five haplotypes of *C. indicum, C. vestitum, C. vestitum* var. *latifolium* and the hybrid between *C. indicum* and *C. vestitum* (Fig. 5). This indicates that either of these species potentially acted as the maternal parent of cultivated chrysanthemums. This is consistent with previous research implicating their involvement (Dai et al., 1998; Dai et al., 2005; Fukai, 2003). However, the specific maternal parental progenitor of modern cultivated chrysanthemums remains elusive. The modern cultivated chrysanthemums were placed in a monophyletic clade that was sister to *C. lavandulifolium*. However, the chloroplast genome of *C. lavandulifolium* has some unique mutations compared with cultivated chrysanthemums. This leads to the hypothesis that the maternal progenitor of modern cultivated chrysanthemums has gone extinct (Ma et al., 2020). However, this requires further evaluation as only one or two whole chloroplast genomes were included for each wild *Chrysanthemum* species and cultivar. This means the diversity of chloroplast genomes was not sufficiently represented. Given the high haplotype diversity of the three taxa and the fact that these haplotypes did not form a monophyletic clade (Fig. 5), we predict that the ultimate maternal progenitor of modern cultivated chrysanthemums may potentially be any of the wild *Chrysanthemum* species, and there may be multiple maternal progenitors for chrysanthemum cultivars.

## Acknowledgements

This work was funded by the National Natural Science Foundation of China (31770230, 31600295 and 31872710), Shandong Provincial Natural Science Foundation, China (ZR2018PC022) and Funds of Shandong ―Double Tops‖ Program (SYL2017XTTD13).

## Tables

**Table S1** Detailed sampling information for natural populations of *C. indicum, C. vestitum* and *C. vestitum* var. *latifolium*.

**Table S2** Details of microsatellite markers used in this study.

**Table S3** Genetic diversity of 14 populations of *C. indicum, C. vestitum* and *C. vestitum* var. *latifolium* based on microsatellite markers.

**Table S4** The chloroplast haplotypes and their variable sites based on *trn*L-*trn*F sequences.

## Figure legends

**Fig. S1.**
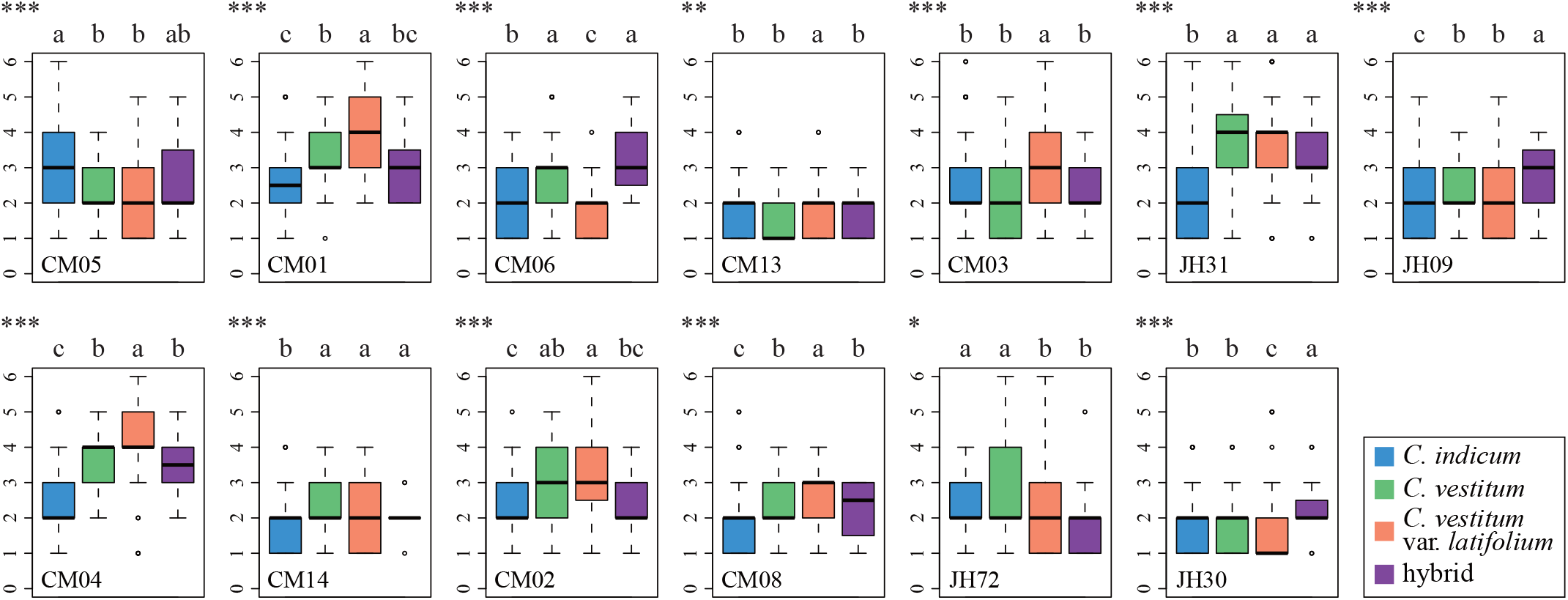
Allele number per individual at each of the 13 microsatellite loci among *C. indicum, C. vestitum, C. vestitum* var. *latifolium* and hybrids. The difference in the number of allele was assessed using Kruskal–Wallis test. ^*^*P* < 0.05, ^**^*P* < 0.01, ^***^*P* < 0.001.

**Fig. S2.**
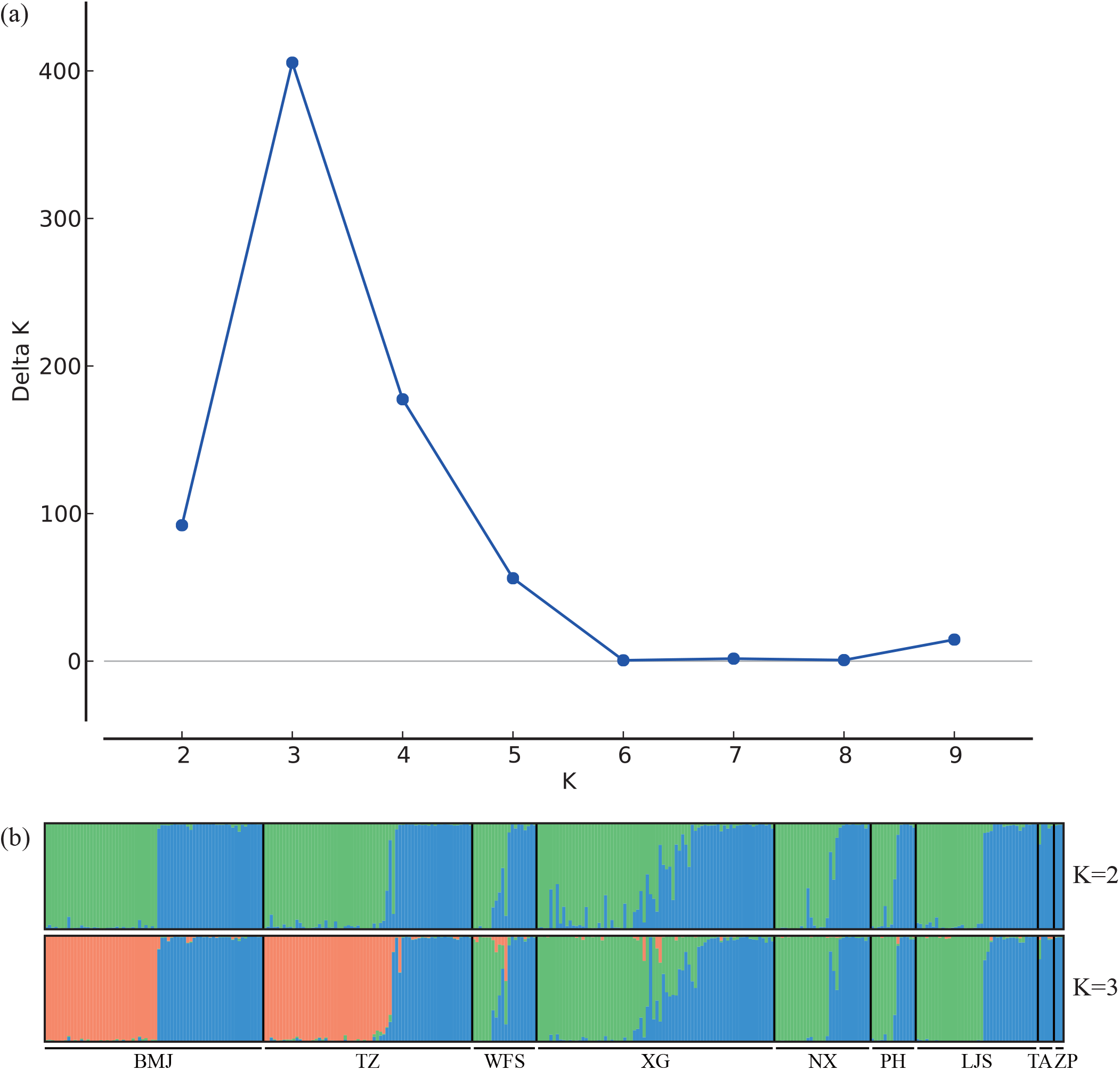
The output of Structure Harvester showing that K = 3 is the optimal value (a) and STRUCTURE results at K = 2 and 3 (b).

